# Feature-specific prediction errors for visual mismatch

**DOI:** 10.1101/447243

**Authors:** Gabor Stefanics, Klaas Enno Stephan, Jakob Heinzle

**Affiliations:** Translational Neuromodeling Unit (TNU), Institute for Biomedical Engineering, University of Zurich & ETH Zurich, Wilfriedstrasse 6, 8032 Zurich, Switzerland; Laboratory for Social and Neural Systems Research, Department of Economics, University of Zurich, Blümlisalpstrasse 10, 8006 Zurich, Switzerland; Max Planck Institute for Metabolism Research, Cologne, Germany

**Keywords:** predictive coding, precision weighted prediction error, color perception, emotion recognition, perception, perceptual inference

## Abstract

Predictive coding (PC) theory posits that our brain employs a predictive model of the environment to infer the causes of its sensory inputs. A fundamental but untested prediction of this theory is that the same stimulus should elicit distinct precision weighted prediction errors (pwPEs) when different (feature-specific) predictions are violated, even in the absence of attention. Here, we tested this hypothesis using functional magnetic resonance imaging (fMRI) and a multi-feature roving visual mismatch paradigm where rare changes in either color (red, green), or emotional expression (happy, fearful) of faces elicited pwPE responses in human participants. Using a computational model of learning and inference, we simulated pwPE and prediction trajectories of a Bayes-optimal observer and used these to analyze changes in blood oxygen level dependent (BOLD) responses to changes in color and emotional expression of faces while participants engaged in a distractor task. Controlling for visual attention by eye-tracking, we found pwPE responses to unexpected color changes in the fusiform gyrus. Conversely, unexpected changes of facial emotions elicited pwPE responses in cortico-thalamo-cerebellar structures associated with emotion and theory of mind processing. Predictions pertaining to emotions activated fusiform, occipital and temporal areas. Our results are consistent with a general role of PC across perception, from low-level to complex and socially relevant object features, and suggest that monitoring of the social environment occurs continuously and automatically, even in the absence of attention.

**Highlights:**

Changes in color or emotion of physically identical faces elicit prediction errors
Prediction errors to such different features arise in distinct neuronal circuits
Predictions pertaining to emotions are represented in multiple cortical areas
Feature-specific prediction errors support predictive coding theories of perception

## Introduction

Predictive coding (PC) postulates that perceptual inference rests on probabilistic (generative) models of the causes of the sensory input (Rao and Ballard, 1999; Friston, 2005; Clark, 2015). The theory emphasizes the active nature of perceptual inference: in contrast to theories that view perception as a reactive, feed-forward analysis of bottom-up sensory information (Hubel and Wiesel, 1965; Riesenhuber and Poggio, 2000), PC regards the brain as actively predicting the sensory signal, based on a hierarchical probabilistic model of the causes of its sensory signals (Egner et al., 2010; Friston, 2010; Lochmann et al., 2012; Bogacz, 2017). According to this theory, perception involves inferring the most likely cause of the sensory signals by integrating incoming sensory information at a given level in the hierarchy with predictions generated at the level above (Rao and Ballard, 1999; Lee and Mumford, 2003; Friston, 2005), where the latter derive from prior information. In this framework a unified perceptual representation of an object involves a set of hierarchical predictions that relate to the object’s different attributes, such as spatiotemporal coordinates but also intrinsic structure. At each hierarchical level, incoming signals from the level below are compared to predictions from the level above, and the ensuing prediction errors (PEs) are passed to the higher level in order to update predictions.

PC thus offers a framework to describe how object representations emerge during hierarchical perceptual inference: segregation and integration of predicted lower-level and more abstract attributes take place in a probabilistic network bound together by passing messages between hierarchical levels that most effectively minimize perceptual PEs (Friston, 2005; Bogacz, 2017). In this framework, unexpected stimuli trigger PE responses which subside as stimuli become predictable, for example through repeated presentation.

PC has become one of the most influential theories of perception, and many of its implications have been confirmed experimentally (e.g., Smith and Muckli, 2010; Wacongne et al., 2011; Kok et al., 2012a,b; Durschmid et al., 2016; Sedley et al., 2016; Ehinger et al., 2017; Gordon et al., 2017; Schwiedrzik and Freiwald, 2017). One central question about the implementation of PC is whether the same physical stimulus elicits separable feature-specific PE responses when distinct predictions about its various attributes exist, regardless whether such attributes are behaviorally relevant. To our knowledge, this has only been studied under attention (Jiang et al., 2016), but not for automatic processing, in the absence of attention and task-relevance. To answer this question, we used a roving standard paradigm (Fig.1A) to systematically manipulate predictions of two attributes of complex stimuli, the color and emotional expression of faces. Based on prior event-related brain potential (ERP) studies, we used a visual mismatch paradigm (for reviews, see Stefanics et al., 2014; Kremlacek et al., 2016) to study brain responses reflecting PEs and model updating processes elicited by unexpected changes in color and facial emotion while participants engaged in a distractor task.

**Fig. 1.**
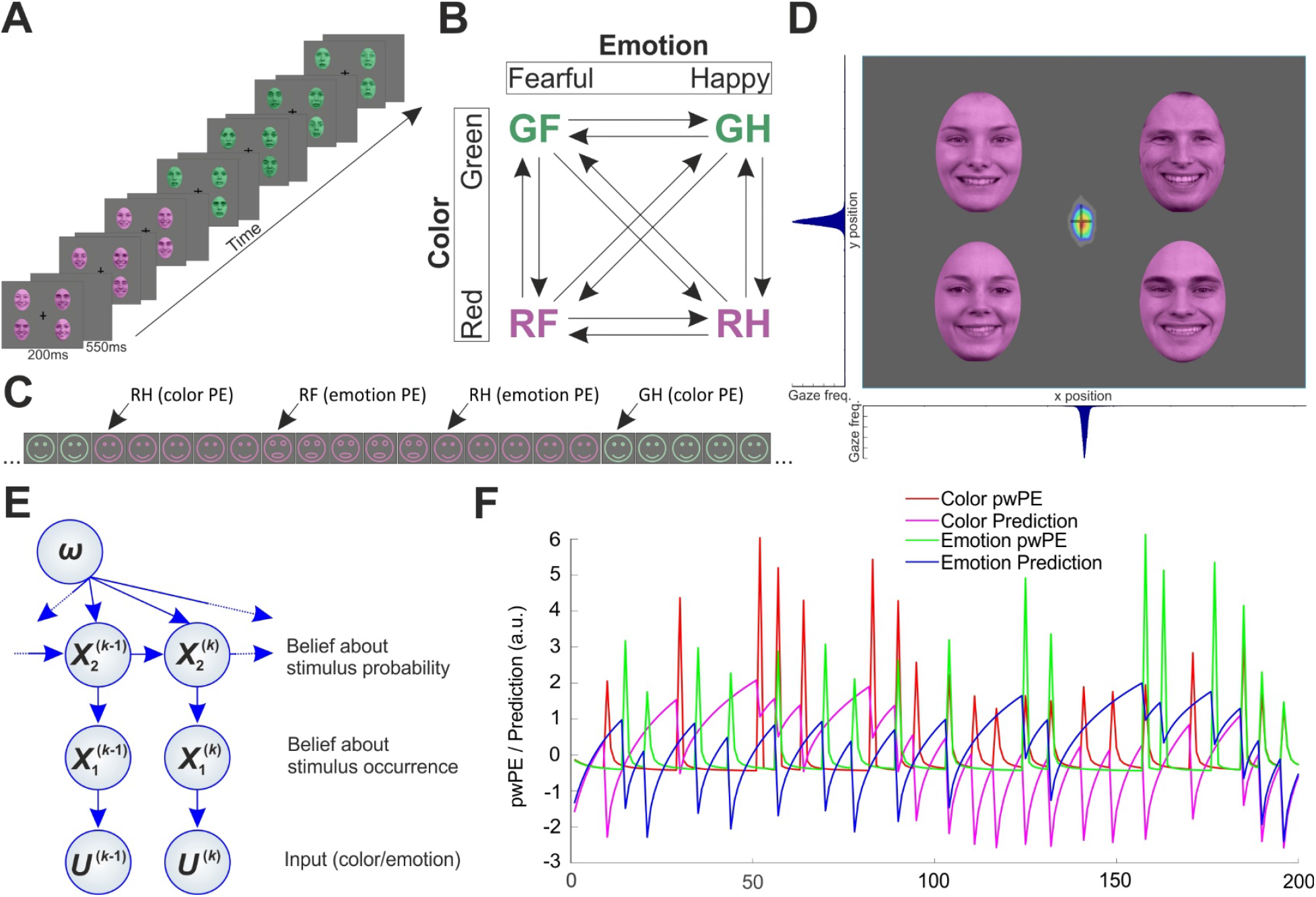
Experimental design and eye-tracking results. A) Four individual photographs of the same color displaying the same facial affect were presented in each stimulus panel for 200ms in a roving standard paradigm. Each panel was followed by an empty grey screen presented for 550ms. The vertical and horizontal lines of the fixation cross occasionally flipped during this interstimulus interval. The subjects’ task was to press a button when the cross flipped. B) Schematic contingency table showing the four equally probable stimulus types (GF: green fearful, GH: green happy, RF: red fearful, RH: red happy faces). After 5-9 presentations each stimulus type was followed by any of the other three types. Arrows indicate transitions with equal overall probability between stimulus types during the experiment. C) Schematic illustration of a stimulus sequence showing transitions between stimulus types. Note physically identical stimuli evoking different PEs depending on expectations established by prior stimulus context. D) Heatmap of normalized gaze position frequency overlaid on a stimulus panel. Warmer colors represent more frequent gaze position. Normalized histograms below and left to the heatmap show the same data projected onto the x and y axis, respectively. Faces were reproduced with permission of the Radboud Faces Database (www.rafd.nl). E) A graphical model of the Hierarchical Gaussian Filter with two levels. F) Model-based pwPE trajectories from one experimental block used as regressors in the GLM.

We used the Hierarchical Gaussian Filter (HGF, Mathys et al., 2011; Mathys et al., 2014) to simulate belief trajectories of an ideal Bayesian observer. The HGF is a computational model that allows for inferring an agent’s beliefs and uncertainty about hidden states of the world that generate sensory information. The model tracks the beliefs of the agent about the probability of each stimulus feature and updates its inference as new information is presented trial-by-trial. The HGF implements a form of PC in the temporal domain and has been used in multiple studies to investigate predictive processes in the brain (e.g., Iglesias et al., 2013; Schwartenbeck et al., 2015; Vossel et al., 2015; Auksztulewicz et al., 2017; Diaconescu et al., 2017; Lawson et al., 2017; Powers et al., 2017; Adams et al., 2018; Katthagen et al., 2018; Stefanics et al., 2018a).

In this paper we used a similar experimental paradigm, computational modeling and analysis approach as in a previous single-trial EEG study that allowed us to study the time course of event-related brain potentials (ERP) to unexpected color and emotion changes associated with pwPEs (Stefanics et al., 2018a). In this previous study, we found that both kind of changes elicited brain responses that were better explained with pwPEs as parametric regressors than regressors encoding categorical stimulus changes in a general linear modeling (GLM) analysis. Here, we used fMRI to identify the brain regions associated with feature-specific predictions and pwPEs to human faces. Critically, our paradigm independently manipulated the color and emotional expression of face stimuli (Fig. 1B, C), allowing us to model predictions and pwPEs to violations of emotion expectations separately from predictions and pwPEs elicited by changes in color. This enabled us to study predictive processes pertaining to low versus high level object features for physically identical stimuli.

## Methods

### Ethics Statement

The experimental protocol was approved by Cantonal Ethics Commission of Zurich (KEK 2010-0327). Written informed consent was obtained from all participants after the procedures and risks were explained. The experiments were conducted in compliance with the Declaration of Helsinki.

### Subjects

Thirty-nine healthy, right-handed subjects participated in this experiment. One subject was excluded due to incomplete data, and three subjects’ data of one scanning day were lost during transfer due to a technical failure. The final sample comprised 35 subjects (mean age=23.06ys, sd=3.02ys, 15 females). All subjects had normal or corrected-to-normal vision.

### Paradigm

Faces were presented in four peripheral quadrants of the screen (Fig. 1A) on a grey background with a fixation cross in the center. Each stimulus panel contained four faces of different identity expressing the same emotion. Stimulus duration was 200ms. The stimuli were presented after an inter-stimulus interval of 550ms during which only the fixation cross was present. A change detection task was presented at the central fixation cross. Roving paradigms have frequently been used to study automatic sensory expectation effects (Haenschel et al., 2005; Garrido et al., 2008; Costa-Faidella et al., 2011; Moran et al., 2013; Auksztulewicz and Friston, 2015; Stefanics et al., 2018a,b). Here, we used a factorially structured multi-feature visual ‘roving standard’ paradigm to elicit PE responses by unexpected changes either in color (red, green), or emotional expression (happy, fearful) of human faces, or both. Importantly, this allowed us to study how brain responses to physically identical stimuli differed, depending on the degree of expectations about color and emotion, respectively. A diagram of the transitions between stimulus types is shown in Fig. 1B.

Images were taken from the Radboud Faces Database (Langner et al., 2010). Ten female and ten male Caucasian models were selected based on their high percentage of agreement on emotion categorization (98% for happy, 92% for fearful faces). A Wilcoxon rank sum test indicated that categorization agreement on the emotional expressions did not differ between happy and fearful faces (Z=-0.63, p=0.53). To control low-level image properties, we equated the luminance and the spatial frequency content of grayscale images of the selected happy and fearful faces using the SHINE toolbox (Willenbockel et al., 2010). The resulting images were used to create the colored stimuli.

### Behavioral task

Similar to previous studies (e.g., Astikainen et al., 2009; Kimura et al., 2012; Müller et al., 2010; Stefanics et al., 2011, 2012, 2018a,b; Kreegipuu et al., 2013; Kuldkepp et al., 2013; Kovács-Bálint et al., 2014; Farkas et al., 2015) we used a behavioral task to engage participants’ attention and thus reduce attentional effects on the processing of face stimuli across participants. The task involved detecting changes in the length of the horizontal and vertical lines of a fixation cross presented in the center of the visual field. At random times, the cross became wider or longer (Fig. 1A), at a rate of 8 flips per minute on average. The cross-flips were unrelated to the changes of the unattended faces. The task was to quickly respond to the cross-flips with a right hand button-press. Reaction times were recorded.

### Eye-tracking

Participants were explicitly asked to fixate at the cross in the center of the screen. To make sure that participants did not direct their overt attention to the face stimuli, we used an Eyelink 1000 eye-tracking system to record gaze position at 250 Hz during the experiment. After removal of intervals immediately before and after, as well as during blinks, heatmap of x-y data points for all subjects were plotted using the EyeMMV toolbox (Krassanakis et al., 2014). A Gaussian filter (SD=3 pixels) was applied to smooth the final image. The heatmap was normalized to have maximum value of 1, and gaze position histograms for x and y coordinates were plotted (Fig.1D).

### Data acquisition and preprocessing

FMRI data was acquired on a Philips Achieva 3 Tesla scanner using an eight channel head-coil (Philips, Best, The Netherlands) at the Laboratory for Social and Neural Systems Research at the University of Zurich. A structural image was acquired for each participant with a T1-weighted MPRAGE sequence: 181 sagittal slices, field of view (FOV): 256 × 256 mm2, Matrix: 256 x 256, resulting in 1 mm^3^ resolution. Functional imaging data was acquired in six experimental blocks. In each block 200 whole-brain images were acquired using a T2*-weighted echo-planar imaging sequence with the following parameters. 42 ascending transverse plane slices with continuous in-plane acquisition (slice thickness: 2.5 mm; in-plane resolution: 3.125 x 3.125 mm; inter-slice gap: 0.6 mm; TR = 2.451 ms; TE = 30 ms; flip angle = 77; field of view = 220 x 220 x 130 mm; SENSE factor = 1.5; EPI factor = 51). We used a 2nd order pencil-beam shimming procedure provided by Philips to reduce field inhomogeneities during the functional scans. All functional images were reconstructed with 3 mm isotropic resolution. Functional data acquisition lasted approximately 1 hour. During fMRI data acquisition, respiratory and cardiac activity was recorded using a breathing belt and an electrocardiogram, respectively.

We used statistical parametric mapping (SPM12, v6470; RRID: SCR_007037; Friston et al., 2007) for fMRI data analysis. First, functional images were slice time corrected, realigned to correct for motion and co-registered with the subject’s own anatomical image. Next, we normalized structural images to MNI space using the unified segmentation approach and applied the same warping to normalize functional images. The functional images were smoothed with a 6 mm full-width at half maximum Gaussian kernel and resampled to 2 mm isotropic resolution. We used RETROICOR (Glover et al., 2000) as implemented in the PhysIO-Toolbox (Kasper et al., 2017) from the open source software TAPAS (http://www.translationalneuromodeling.org/tapas) to create confound regressors for cardiac pulsations, respiration, and cardio-respiratory interactions. These confound regressors were entered into the general linear model (GLM; see below). The data and code used in this study are available from the corresponding author, upon reasonable request.

### Modeling belief trajectories

In order to include parametric regressors of precision weighted prediction errors (pwPE) in the GLM, we simulated trajectories of belief update in a generative model of perceptual inference, the Hierarchical Gaussian Filter (HGF; Mathys et al., 2011; 2014). We followed the approach described in details in Stefanics et al. (2018a) using the HGF toolbox version v2.2 contained in TAPAS (http://www.translationalneuromodeling.org/tapas). Briefly, we simulated the perceptual model of a two-level HGF for the input traces given by the two features of the face stimuli: color (red vs. green) and emotion (fearful vs. happy). Inversion of the HGF (Fig. 1E) infers the hidden states (*x*) of the world that generate the sensory input (*u*). The belief states are updated after each trial following a generic update rule: The posterior mean 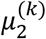 of state *x*_2_ at trial k changes its value according to a precision-weighted PE 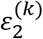, where the precision-weighting changes trial by trial and can be regarded as dynamic learning rate:

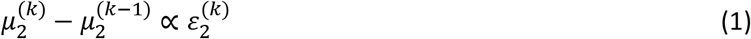

Note that the sigmoid transform of the tendency 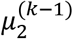 constitutes the prediction (probability of observing an input 1 on trial k), while 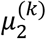 is the tendency after it was updated according to the input on trial k. Here, we refer to 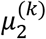 as prediction. For comparison, classical associative and reinforcement learning models (e.g., Rescorla and Wagner, 1972) follow a similar form but use a fixed learning rate:

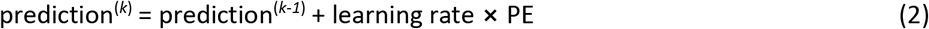

For the simulations we assumed that color and emotion were processed by two separate, independent HGFs. However, we considered an interaction between color and emotion PEs within a GLM. Investigating possible interactions at the level of the perceptual model of the HGF would require establishing a novel version of the HGF that incorporates interactions between hidden beliefs, which was beyond the scope of our current study. We estimated the parameters of the model assuming an ideal Bayes-optimal observer (Mathys et al., 2011) that minimizes surprise of the incoming input stream. Figure 1F displays example traces of the absolute value of *μ*_2_ and *ε*_2_ which entered the GLM as described below.

### General linear model analysis

The fMRI data was analyzed with two separate GLMs. One GLM included the gradually changing (absolute) pwPEs and “prediction strength” given by the absolute value of *μ*_2_ derived from the HGF as modulatory regressors while the other GLM incorporated a regressor representing categorical stimulus change. The latter served for comparison, implementing a simpler alternative than PC, i.e., change detection (CD; see Lieder et al., 2013). For the GLM based on the CD model, we included stick functions as parametric modulators for each stimulus on those trials when a change occurred in the stimulus sequence. The GLMs were estimated for each participant individually. The pwPE and prediction strength as well as the CD modulatory regressors were computed separately for color and emotion. In addition the GLM included modulatory regressors for red vs. green and happy vs. fearful, respectively. Hence, for each run of the experiment the design matrix included the following experimental regressors: i) a main regressor for the onset of each stimulus display, ii) two modulatory regressors encoding color (red = −1, green = 1) and emotion (happy = −1, fearful = 1), respectively, and iii) two modulatory regressors with the absolute pwPE (or CD) for color and emotion, and iv) two modulatory regressors with the absolute value of the tendency (|*μ*_2_|) for color and emotion (only, in the case of the HGF based model). The modulatory regressors were mean centered and normalized to unit variance. In addition to these regressors of interest, button presses to cross-flips of the visual attention task were also included in the model. All regressors were convolved with a canonical hemodynamic response function (HRF). Movement regressors and physiological confounds were included in the first level GLM (Kasper et al., 2017) which was estimated for each participant individually. Please note that the sign of colors and emotions in ii) was arbitrarily chosen. Finally, in order to assess whether there was any interaction between color and emotion PEs, we fitted an additional GLM, where we included the Hadamard (element-wise) product of the color and emotion pwPE as an additional regressor.

On the group level, we used F-tests to find regions whose response showed significant correlation with pwPE or stick regressors. The resulting statistical parametric maps (SPM) were family-wise error (FWE) corrected at the cluster level (p<0.05) with a cluster defining threshold (CDT) of p<0.001 (Woo et al., 2014; Flandin and Friston, 2017). We used probabilistic anatomical labels and cytoarchitectonic maps in the SPM Anatomy toolbox (v2.2c; RRID: SCR_013273, Eickhoff et al., 2005) to identify the anatomical areas/structures where we observed significant effects. We summarize activations in terms of anatomical labeling by reporting all local maxima within each cluster in Table 1. This provides an overview over the activations in terms of commonly used anatomical labels.

**Table 1.**
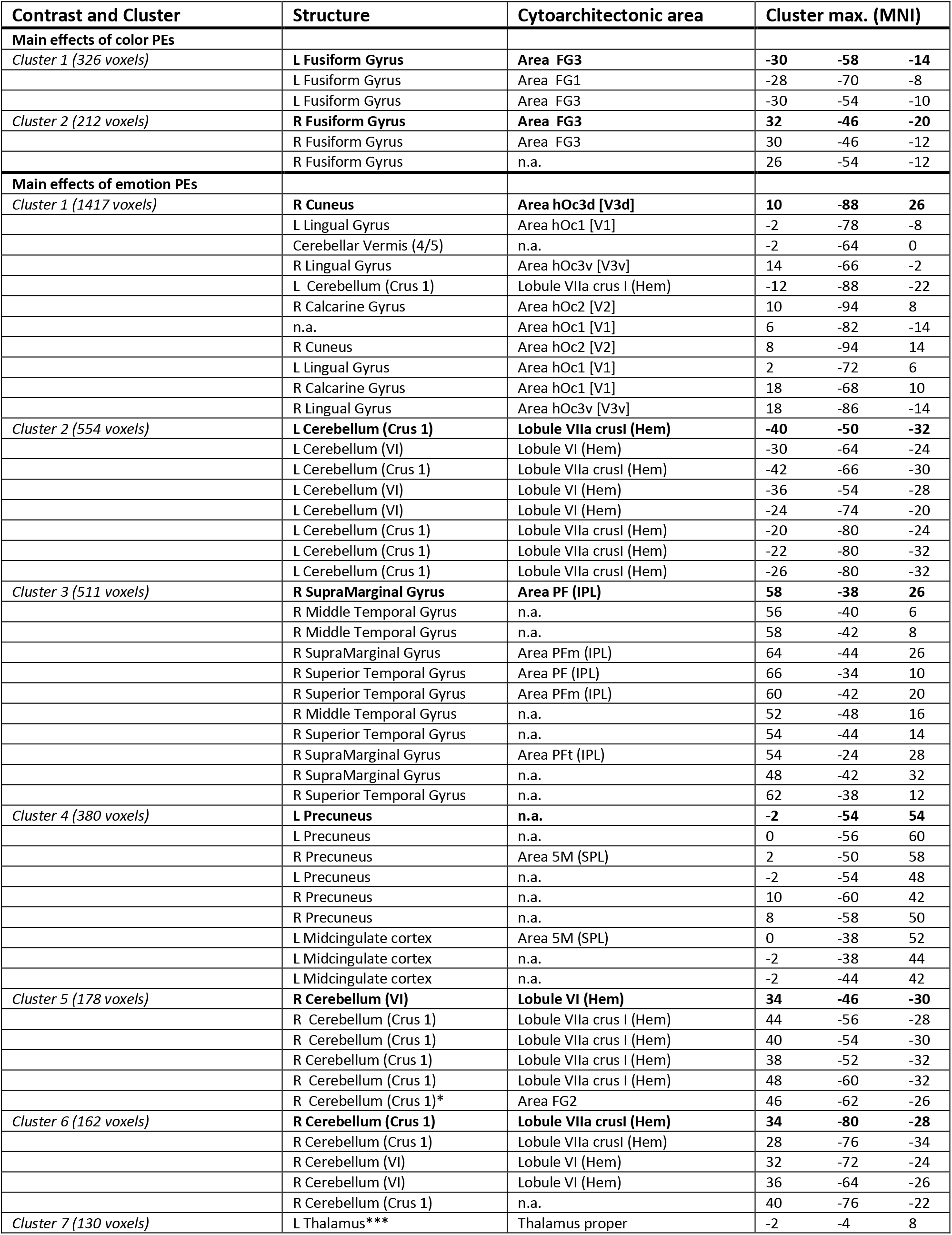
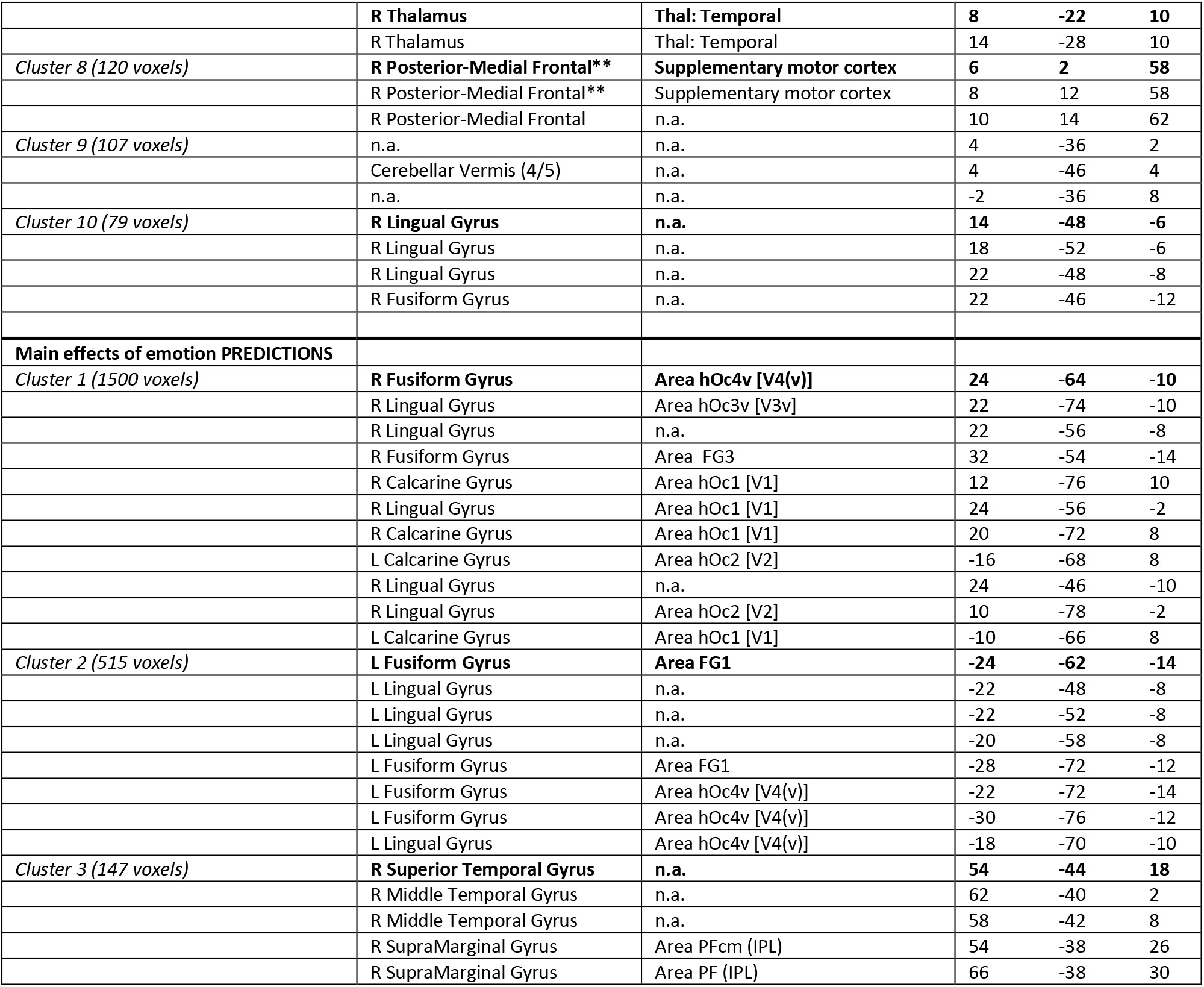
Assignment of activations to anatomical and cytoarchitectonic regions (Anatomy Toolbox, v2.2c). In order to characterize the anatomical locations of the cluster we report maxima within the clusters and their assignment to anatomical regions. If a maximum lies within a particular region, this means that the cluster extends into that anatomical region, but does not imply that the entire region is activated or that the entire cluster lies within that anatomical region. Whole brain analyses on the cluster level p<0.05 (FWE-corrected) with a cluster defining threshold of p<0.001. Contrast estimates from structures in bold font are plotted in Figures 2–4. n.a.: these maxima were not assigned to any region. *The anatomy toolbox labelled this maximum as Cerebellum but assigned it to the fusiform area FG2. **The anatomy toolbox labelled this maximum as Posterior-Medial Frontal cortex but did not assign it. The anatomical label was corrected to Supplementary motor cortex based on Neuromorphometrics labelling in SPM. ***The anatomy toolbox did not label this maximum. The anatomical label of left Thalamus was added based on Neuromorphometrics labelling in SPM.

| Contrast and Cluster | Structure | Cytoarchitectonic area | Cluster max. (MNI) |  |  |
| --- | --- | --- | --- | --- | --- |
| **Main effects of color PEs** |  |  |  |  |  |
| Cluster 1 (326 voxels) | **L Fusiform Gyrus** | **Area FG3** | **-30** | **-58** | **-14** |
|  | L Fusiform Gyrus | Area FG1 | -28 | -70 | -8 |
|  | L Fusiform Gyrus | Area FG3 | -30 | -54 | -10 |
| Cluster 2 (212 voxels) | **R Fusiform Gyrus** | **Area FG3** | **32** | **-46** | **-20** |
|  | R Fusiform Gyrus | Area FG3 | 30 | -46 | -12 |
|  | R Fusiform Gyrus | n.a. | 26 | -54 | -12 |
| **Main effects of emotion PEs** |  |  |  |  |  |
| Cluster 1 (1417 voxels) | **R Cuneus** | **Area hOc3d [V3d]** | **10** | **-88** | **26** |
|  | L Lingual Gyrus | Area hOc1 [V1] | -2 | -78 | -8 |
|  | Cerebellar Vermis (4/5) | n.a. | -2 | -64 | 0 |
|  | R Lingual Gyrus | Area hOc3v [V3v] | 14 | -66 | -2 |
|  | L Cerebellum (Crus 1) | Lobule VIIa crus I (Hem) | -12 | -88 | -22 |
|  | R Calcarine Gyrus | Area hOc2 [V2] | 10 | -94 | 8 |
|  | n.a. | Area hOc1 [V1] | 6 | -82 | -14 |
|  | R Cuneus | Area hOc2 [V2] | 8 | -94 | 14 |
|  | L Lingual Gyrus | Area hOc1 [V1] | 2 | -72 | 6 |
|  | R Calcarine Gyrus | Area hOc1 [V1] | 18 | -68 | 10 |
|  | R Lingual Gyrus | Area hOc3v [V3v] | 18 | -86 | -14 |
| Cluster 2 (554 voxels) | **L Cerebellum (Crus 1)** | **Lobule VIIa crusI (Hem)** | **-40** | **-50** | **-32** |
|  | L Cerebellum (VI) | Lobule VI (Hem) | -30 | -64 | -24 |
|  | L Cerebellum (Crus 1) | Lobule VIIa crusI (Hem) | -42 | -66 | -30 |
|  | L Cerebellum (VI) | Lobule VI (Hem) | -36 | -54 | -28 |
|  | L Cerebellum (VI) | Lobule VI (Hem) | -24 | -74 | -20 |
|  | L Cerebellum (Crus 1) | Lobule VIIa crusI (Hem) | -20 | -80 | -24 |
|  | L Cerebellum (Crus 1) | Lobule VIIa crusI (Hem) | -22 | -80 | -32 |
|  | L Cerebellum (Crus 1) | Lobule VIIa crusI (Hem) | -26 | -80 | -32 |
| Cluster 3 (511 voxels) | **R SupraMarginal Gyrus** | **Area PF (IPL)** | **58** | **-38** | **26** |
|  | R Middle Temporal Gyrus | n.a. | 56 | -40 | 6 |
|  | R Middle Temporal Gyrus | n.a. | 58 | -42 | 8 |
|  | R SupraMarginal Gyrus | Area PFm (IPL) | 64 | -44 | 26 |
|  | R Superior Temporal Gyrus | Area PF (IPL) | 66 | -34 | 10 |
|  | R Superior Temporal Gyrus | Area PFm (IPL) | 60 | -42 | 20 |
|  | R Middle Temporal Gyrus | n.a. | 52 | -48 | 16 |
|  | R Superior Temporal Gyrus | n.a. | 54 | -44 | 14 |
|  | R SupraMarginal Gyrus | Area PFt (IPL) | 54 | -24 | 28 |
|  | R SupraMarginal Gyrus | n.a. | 48 | -42 | 32 |
|  | R Superior Temporal Gyrus | n.a. | 62 | -38 | 12 |
| Cluster 4 (380 voxels) | **L Precuneus** | **n.a.** | **-2** | **-54** | **54** |
|  | L Precuneus | n.a. | 0 | -56 | 60 |
|  | R Precuneus | Area 5M (SPL) | 2 | -50 | 58 |
|  | L Precuneus | n.a. | -2 | -54 | 48 |
|  | R Precuneus | n.a. | 10 | -60 | 42 |
|  | R Precuneus | n.a. | 8 | -58 | 50 |
|  | L Midcingulate cortex | Area 5M (SPL) | 0 | -38 | 52 |
|  | L Midcingulate cortex | n.a. | -2 | -38 | 44 |
|  | L Midcingulate cortex | n.a. | -2 | -44 | 42 |
| Cluster 5 (178 voxels) | **R Cerebellum (VI)** | **Lobule VI (Hem)** | **34** | **-46** | **-30** |
|  | R Cerebellum (Crus 1) | Lobule VIIa crus I (Hem) | 44 | -56 | -28 |
|  | R Cerebellum (Crus 1) | Lobule VIIa crus I (Hem) | 40 | -54 | -30 |
|  | R Cerebellum (Crus 1) | Lobule VIIa crus I (Hem) | 38 | -52 | -32 |
|  | R Cerebellum (Crus 1) | Lobule VIIa crus I (Hem) | 48 | -60 | -32 |
|  | R Cerebellum (Crus 1)* | Area FG2 | 46 | -62 | -26 |
| Cluster 6 (162 voxels) | **R Cerebellum (Crus 1)** | **Lobule VIIa crusI (Hem)** | **34** | **-80** | **-28** |
|  | R Cerebellum (Crus 1) | Lobule VIIa crusI (Hem) | 28 | -76 | -34 |
|  | R Cerebellum (VI) | Lobule VI (Hem) | 32 | -72 | -24 |
|  | R Cerebellum (VI) | Lobule VI (Hem) | 36 | -64 | -26 |
|  | R Cerebellum (Crus 1) | n.a. | 40 | -76 | -22 |
| Cluster 7 (130 voxels) | L Thalamus*** | Thalamus proper | -2 | -4 | 8 |
|  | **R Thalamus** | **Thal: Temporal** | **8** | **-22** | **10** |
|  | R Thalamus | Thal: Temporal | 14 | -28 | 10 |
| *Cluster 8 (120 voxels)* | **R Posterior-Medial Frontal\*\*** | **Supplementary motor cortex** | **6** | **2** | **58** |
|  | R Posterior-Medial Frontal** | Supplementary motor cortex | 8 | 12 | 58 |
|  | R Posterior-Medial Frontal | n.a. | 10 | 14 | 62 |
| *Cluster 9 (107 voxels)* | n.a. | n.a. | 4 | -36 | 2 |
|  | Cerebellar Vermis (4/5) | n.a. | 4 | -46 | 4 |
|  | n.a. | n.a. | -2 | -36 | 8 |
| *Cluster 10 (79 voxels)* | **R Lingual Gyrus** | **n.a.** | **14** | **-48** | **-6** |
|  | R Lingual Gyrus | n.a. | 18 | -52 | -6 |
|  | R Lingual Gyrus | n.a. | 22 | -48 | -8 |
|  | R Fusiform Gyrus | n.a. | 22 | -46 | -12 |
| **Main effects of emotion PREDICTIONS** |  |  |  |  |  |
| *Cluster 1 (1500 voxels)* | **R Fusiform Gyrus** | **Area hOc4v [V4(v)]** | **24** | **-64** | **-10** |
|  | R Lingual Gyrus | Area hOc3v [V3v] | 22 | -74 | -10 |
|  | R Lingual Gyrus | n.a. | 22 | -56 | -8 |
|  | R Fusiform Gyrus | Area FG3 | 32 | -54 | -14 |
|  | R Calcarine Gyrus | Area hOc1 [V1] | 12 | -76 | 10 |
|  | R Lingual Gyrus | Area hOc1 [V1] | 24 | -56 | -2 |
|  | R Calcarine Gyrus | Area hOc1 [V1] | 20 | -72 | 8 |
|  | L Calcarine Gyrus | Area hOc2 [V2] | -16 | -68 | 8 |
|  | R Lingual Gyrus | n.a. | 24 | -46 | -10 |
|  | R Lingual Gyrus | Area hOc2 [V2] | 10 | -78 | -2 |
|  | L Calcarine Gyrus | Area hOc1 [V1] | -10 | -66 | 8 |
| *Cluster 2 (515 voxels)* | **L Fusiform Gyrus** | **Area FG1** | **-24** | **-62** | **-14** |
|  | L Lingual Gyrus | n.a. | -22 | -48 | -8 |
|  | L Lingual Gyrus | n.a. | -22 | -52 | -8 |
|  | L Lingual Gyrus | n.a. | -20 | -58 | -8 |
|  | L Fusiform Gyrus | Area FG1 | -28 | -72 | -12 |
|  | L Fusiform Gyrus | Area hOc4v [V4(v)] | -22 | -72 | -14 |
|  | L Fusiform Gyrus | Area hOc4v [V4(v)] | -30 | -76 | -12 |
|  | L Lingual Gyrus | Area hOc4v [V4(v)] | -18 | -70 | -10 |
| *Cluster 3 (147 voxels)* | **R Superior Temporal Gyrus** | **n.a.** | **54** | **-44** | **18** |
|  | R Middle Temporal Gyrus | n.a. | 62 | -40 | 2 |
|  | R Middle Temporal Gyrus | n.a. | 58 | -42 | 8 |
|  | R SupraMarginal Gyrus | Area PFcm (IPL) | 54 | -38 | 26 |
|  | R SupraMarginal Gyrus | Area PF (IPL) | 66 | -38 | 30 |

## Results

### Fixation and behavioral responses

Gaze position data (Fig. 1D) confirmed that participants complied with task instructions and fixated the central fixation cross throughout the task. Thus, participants engaged in the detection task and were not overtly attending the faces. Mean reaction time to cross-flips was 484ms (standard deviation: SD=106.9ms), and mean hit rate was 78% (SD=7.34%).

### First-level GLMs

We fitted two GLMs on the single-subject level, incorporating parametric regressors that represented two hypotheses about the decay of pwPE/prediction responses following a change in color of emotional expression of the faces. Similar to the model comparison procedure described in our previous study, our original aim was to create a functionally defined mask of significant voxels showing PE responses under both models at the group level (Stefanics et al., 2018a). However, while similar activation clusters were obtained using the pwPE/prediction and CD regressors to color changes, significant clusters to changes in emotion were only found using the pwPE/prediction regressors. In other words, the beta estimates obtained using CD were not consistent enough across subjects to yield significant activation clusters at the group level. The lack of significant group-level results for the CD regressors prevented us from creating an unbiased mask comprising significant voxels for color and emotion (“logical AND” conjunction). Furthermore, the additional analysis which included the interaction (product) of color and emotion pwPE did not reveal any evidence for an interaction between the two. We thus restrict ourselves to reporting the results obtained at the group-level analysis using the HGF-based pwPE/prediction model.

### Effect of color pwPE

A whole-brain analysis of color changes showed significant activation for color pwPE in fusiform areas (Fig. 2A). Post hoc inspection of the contrast estimates (Fig. 2B) revealed an increased response to pwPE. Predictions pertaining to color did not yield significant activations. Detailed information about anatomical labels, cluster size, and MNI coordinates for the maxima of significant voxel clusters are listed in Table 1.

**Fig. 2.**
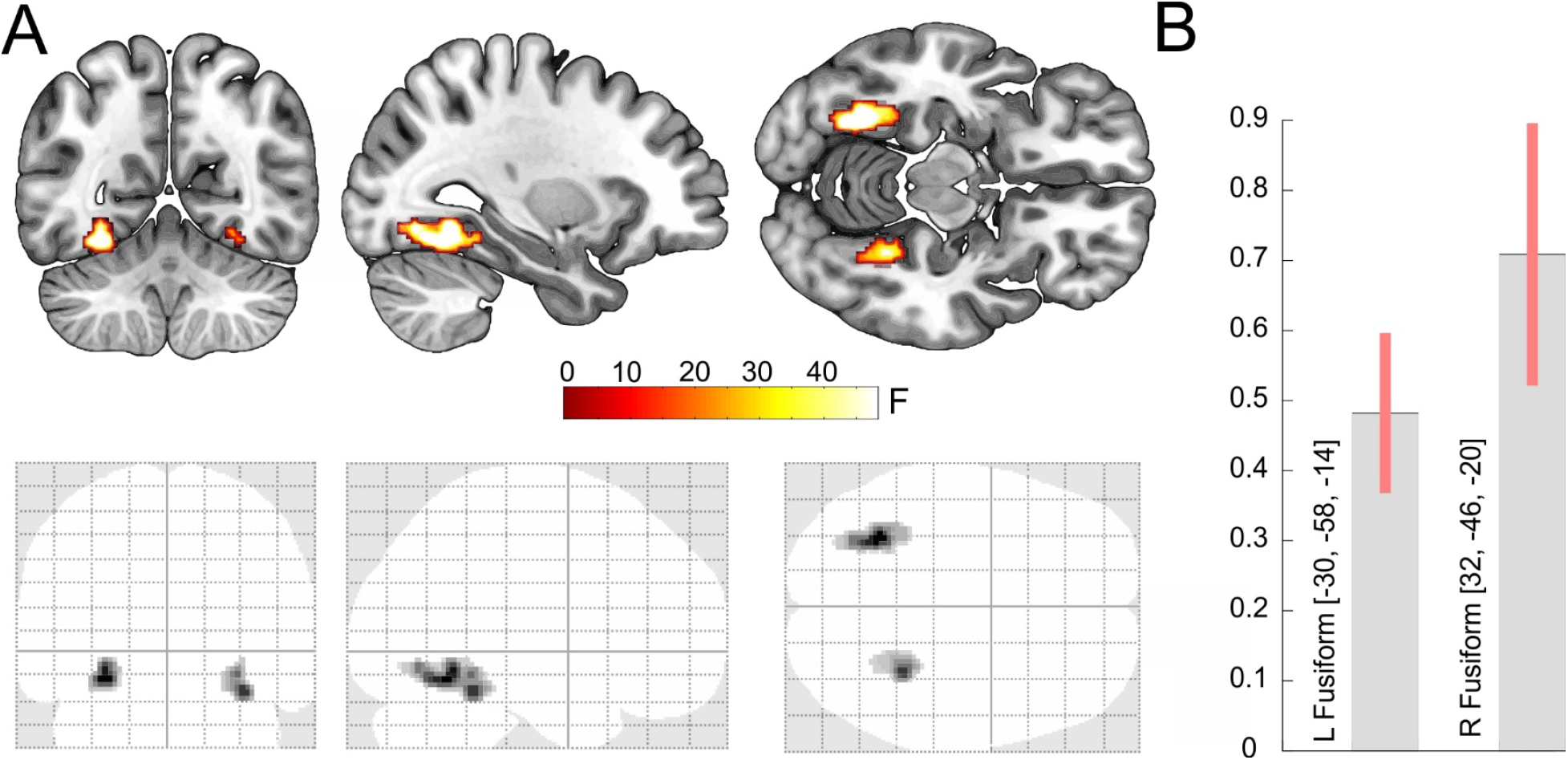
Effect of color pwPE. A) Top: Activation map (p<0.05 cluster-level whole-brain FWE corrected, with a CDT of p<0.001) overlaid onto the MNI152 standard-space T1-weighted average structural template. Slices show activation in the left fusiform gyrus (MNI-coordinates: [−30 −58 −14]. Bottom: Glass brain showing the results of the F-test. B) Contrast estimates (arbitrary units) for color pwPEs in the left and right fusiform gyrus. Bars indicate 90% C.I.

### Effect of emotion prediction and pwPE

A whole-brain analysis of emotion PEs showed significant effects in bilateral cerebellum, cuneus, lingual gyrus, precuneus, thalamus, and right supramarginal gyrus (extending into superior and middle temporal gyrus) as well as right posterior medial frontal cortex (Fig. 3A).

**Fig. 3.**
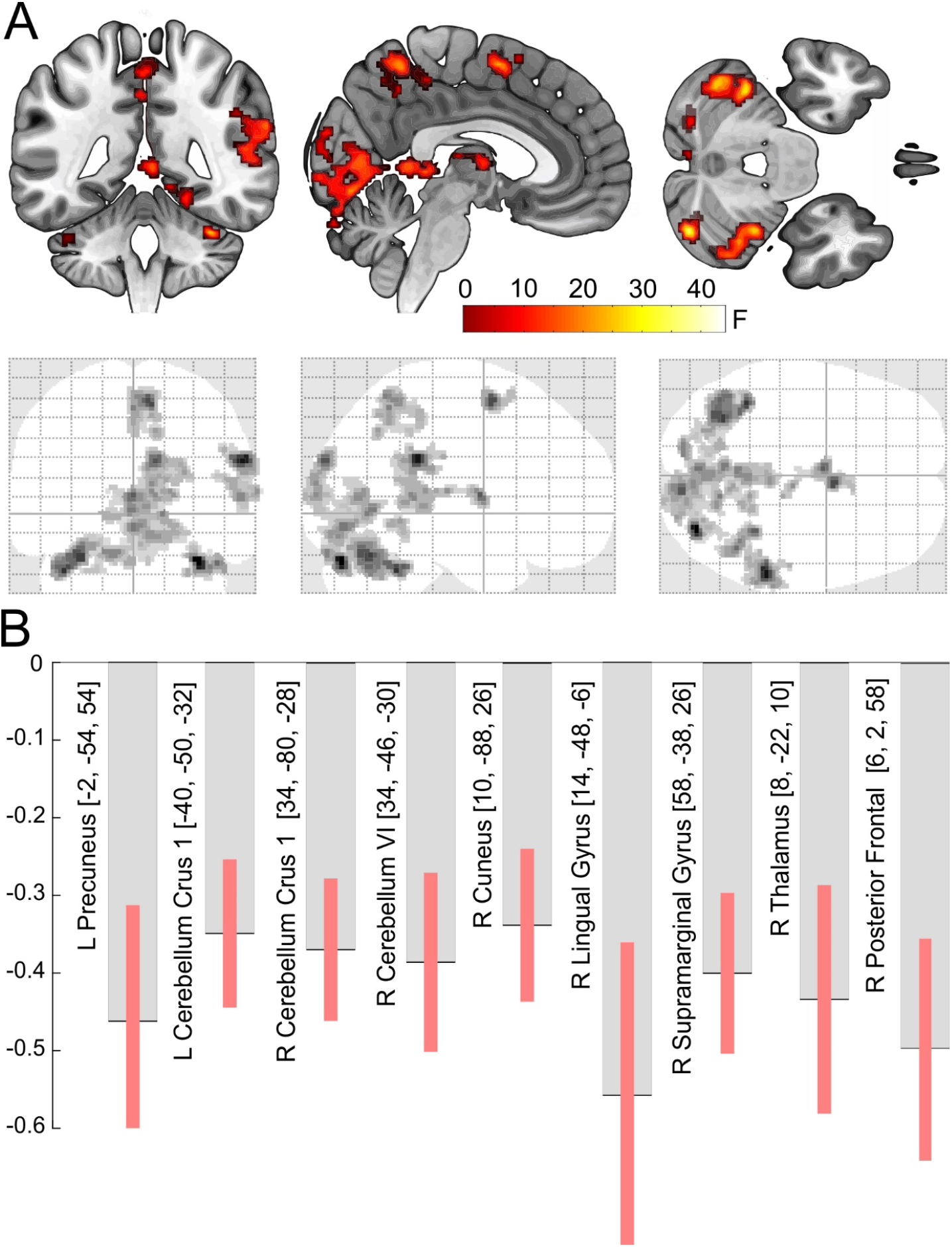
Main effect of emotion pwPE. A) Top: Activation map (p<0.05 cluster-level whole-brain FWE corrected, with a CDT of p<0.001) overlaid onto the MNI152 standard-space T1-weighted average structural template. Slices show activations at coordinates [4, −45, −31] cutting through the right anterior precuneus. Bottom: Glass brain showing the results of the F-test (whole-brain FWE cluster-level corrected at p<0.05, with a cluster-defining threshold of p<0.001). B) Contrast estimates (arbitrary units) for the emotion pwPEs in the left and right cerebellum, left precuneus, right cuneus, lingual and supramarginal gyrus, thalamus, and posterior frontal cortex. Bars indicate 90% C.I. Note that bar plots are shown for illustration only. Statistical significance was assessed at the whole-brain level described above.

We found significant activations pertaining to emotion predictions in three cortical clusters: two in bilateral fusiform gyri (extending into lingual gyri) and one in right the superior and middle temporal cortex (Fig. 4A). A post hoc analysis of the contrast estimates in these regions revealed that all areas showed a negative effect of emotion pwPEs (Fig. 3B) and predictions (Fig. 4B).

**Fig. 4.**
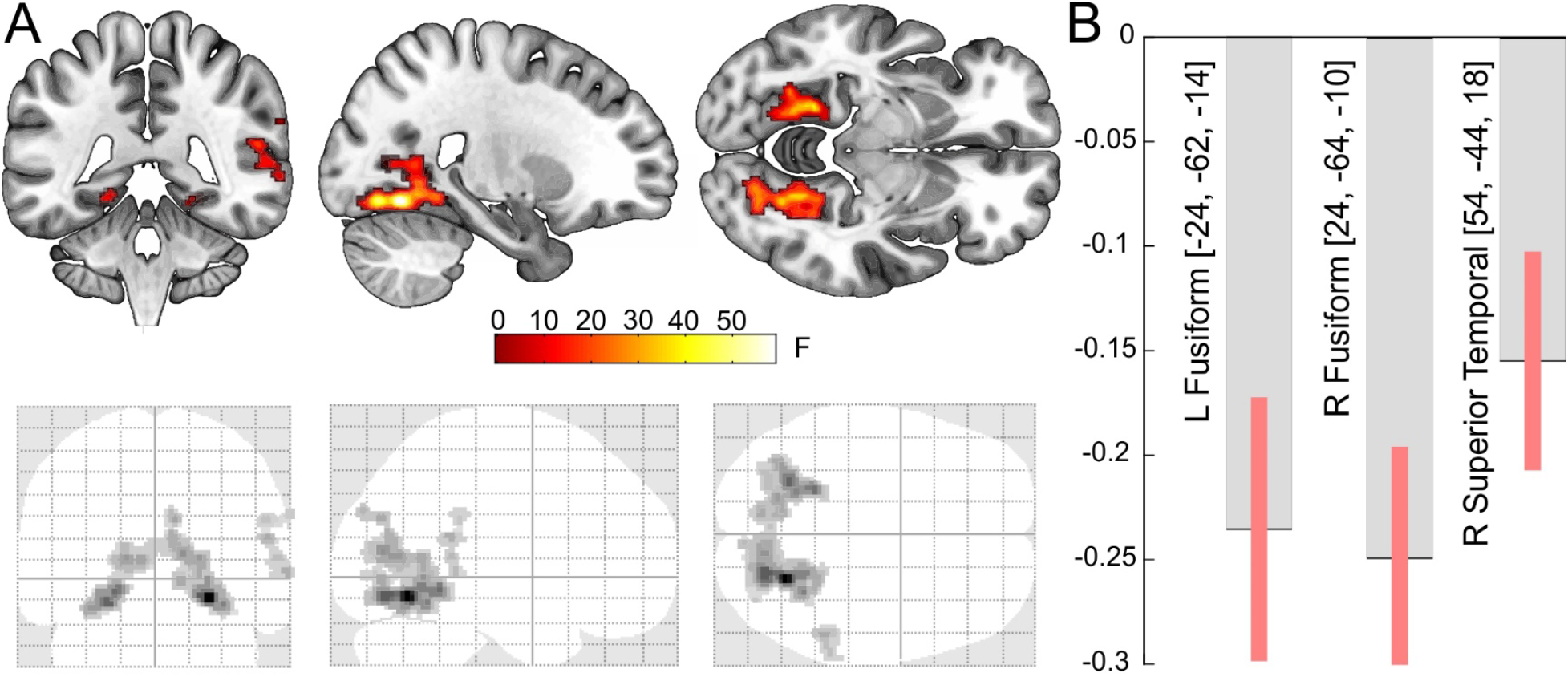
Main effect of emotion prediction. A) Top: Activation map (p<0.05 cluster-level whole-brain FWE corrected, with a CDT of p<0.001) overlaid onto the MNI152 standard-space T1-weighted average structural template. Slices show activations at coordinates [24, −42, −8] cutting through the right fusiform gyrus. Bottom: Glass brain showing the results of the F-test (whole-brain FWE cluster-level corrected at p<0.05, with a cluster-defining threshold of p<0.001). B) Contrast estimates (arbitrary units) for the emotion prediction in the left and right fusiform gyrus, and right superior temporal gyrus. Bars indicate 90% C.I. Note that bar plots are shown for illustration only. Statistical significance was assessed at the whole-brain level described above.

## Discussion

We used the Hierarchical Gaussian Filter, a computational model for learning and inference, to simulate belief trajectories of an ideal Bayesian observer presented with a sequence of face stimuli. The trial by trial update of internal hidden belief states in the HGF relies on precision weighted prediction errors. Traces of predictions and pwPEs pertaining to color and emotional expression of faces served as regressors in a GLM which yielded brain structures where activation showed a significant relationship to those computational quantities. We manipulated sensory expectations towards color and emotional expression of faces independently. Crucially, emotion and color pwPEs/predictions were evoked by physically identical stimuli; only the specific expectation (statistical regularity) that was violated on any given trial, differed between the two conditions. While our previous EEG study reported the scalp distribution and time-course of pwPE responses (Stefanics et al., 2018a), here we used fMRI to find BOLD correlates of pwPEs and predictions in generator structures. We found BOLD correlates of pwPEs to color changes in bilateral fusiform gyrus, whereas pwPEs to changes of emotional expressions activated a different set of areas including the bilateral cerebellum, lingual gyrus, precuneus, thalamus, and right supramarginal gyrus as well as right posterior medial frontal cortex. We observed activations pertaining to emotion predictions in bilateral fusiform and the right supramarginal gyrus (Fig. 5A).

**Fig. 5.**
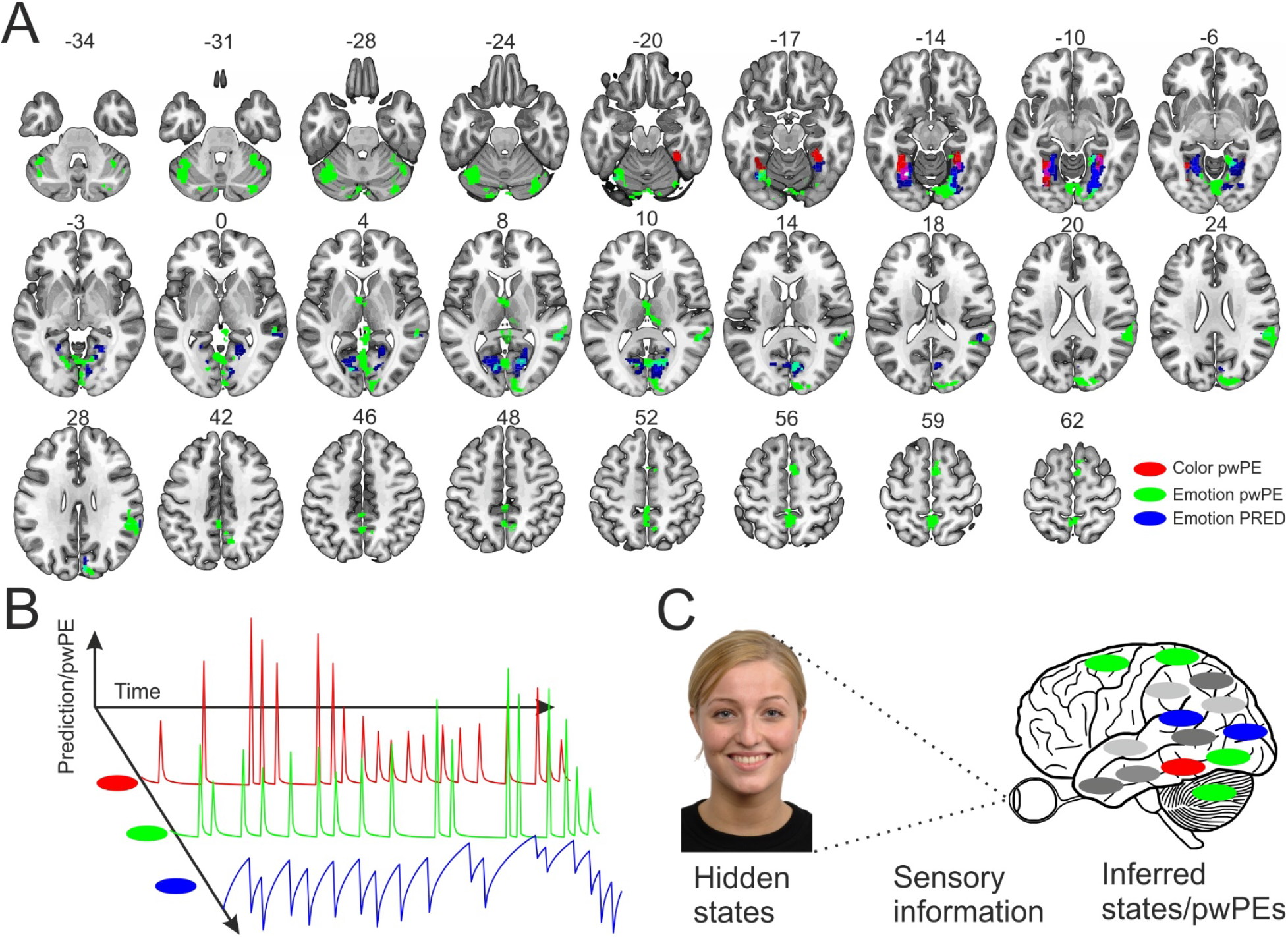
Overview of the results and PC framework for perceptual prediction errors. A) Colored areas mark main clusters related to color pwPEs (red), and emotion pwPEs (green). Note the dissociation of PEs for color and emotion changes. B) pwPE- and prediction-related activations for different sensory features arise and are updated, respectively, during Bayesian inference as properties of the hidden states that cause the sensory information dynamically change over time. Prediction and pwPEs to color and emotion are marked as in A), additional features are marked with grey. C) Schematic depicting functional segregation in the nervous system, as distinct features of the world are inferred and predicted by distinct neural structures specializing in the given features. Image of a model used in our study reproduced with permission of the Radboud Faces Database (Langner et al., 2010).

According to recent hierarchical formulations of PC (Friston, 2005), creating and maintaining our internal model of the world is a process during which predictive object representations about the likely properties of the hidden objects are updated using precision-weighted PEs (e.g., Moran et al., 2013; Stefanics et al., 2018a) that signals mismatch between the expectations based on prior information and the current sensory data (Fig. 5B). In the present study, the demonstration of activations correlated to pwPE in ventral visual areas as well as in emotion processing structures suggests a role for PC in color and emotion perception. Importantly, we manipulated stimulus sequences to induce automatic expectations about the occurrence of different stimulus features, using the same faces to elicit distinct emotion and color pwPEs. In line with our hypothesis, color and emotion pwPEs were reflected by activity in brain structures known to be dedicated to color and emotion processing. A hypothetical generalization of our results is shown in Fig. 5C, which illustrates functional segregation of inferring hidden causes of sensory information for different features, including color and emotional expression of faces.

Here, we studied predictions and pwPEs to unattended and task-irrelevant stimuli. We used a primary task independent of the facial stimuli to ensure that participants did not attend to the faces and verified their attentional focus by eye-tracking. Thus, predictions and pwPEs were elicited under an automatic recognition processes and minimized confounding variations in attentional contributions.

It is important to note that due to the lack of significant group level results for the emotion stick regressor, we were not able to directly compare models using the approach presented in Stefanics et al. (2018a). Hence, we could not use model comparison to assess whether the pwPE or the stick regressor traces provided the formally better model. Notably, the latter are equivalent to a very quick adaptation without precision weighting. However, the second level results suggest that the representation of pwPEs is more consistent across subjects, leading to a significant group effect. In addition, while we use computational quantities to model neural activity in the GLM, our method (fMRI) does not allow us to make a direct statement about the neuronal implementation, e.g., neuronal fatigue, suppressive effects in single neurons, or network effects (e.g., Solomon and Kohn; 2014, Stefanics et al., 2016). Based on the current analysis it is not possible to reject some form of adaptation (e.g., fatigue) as a potential mechanism as opposed to a more general model based on hierarchical Bayesian inference. Thus, adaptation could be an alternative explanation of our findings.

To our knowledge this is the first fMRI study using a Bayesian observer model to describe automatic predictions and pwPEs to violations of expectations to different features of the same objects, in the absence of focal attention and task-relevance. Both expectation based on stimulus probability and attention based on task-relevance have been suggested to modulate sensory PEs (e.g., Summerfield and de Lange, 2014; Auksztulewicz and Friston, 2015; Auksztulewicz et al., 2017). Attentional effects have been suggested to increase synaptic gain of PE coding neurons (Kok et al., 2012; Wyart et al., 2012; Jiang et al., 2013; Vossel et al., 2014; Auksztulewicz and Friston, 2015), whereas expectation effects manifest in reduced neuronal responses (Grotheer and Kovács, 2015; Auksztulewicz and Friston, 2016; Stefanics et al., 2018a,b). Recent formulations of PC suggest that attention serves to optimize precision estimates of specific PEs. By increasing the weight that is put on PEs, the role of attention is to influence subsequent inference and learning (Friston, 2009; Feldman and Friston, 2010; den Ouden et al., 2012; Parr and Friston, 2018). Furthermore, a previous study also found that PEs spread across object features in the visual cortex (Jiang et al., 2016). Here, we extend these previous findings by showing that (i) pwPEs can also be elicited in spatially remote neural structures that specialize in the processing of distinct stimulus attributes and (ii) in the absence of attention. Notably, Jiang et al. (2016) studied PEs to attended and task-relevant random dot stimuli, while in our study face stimuli were task-irrelevant and not attended, as verified by eye tracking. The differences between our current and their results suggest that the role of focal attention in perception might not only be to enhance but also spread PEs across features at the object level (Jiang et al., 2016) which is in line with the feature-integration theory of attention (Treisman and Gelade, 1980). Thus, while the visual system likely represents statistical relationships across features and automatically structures them into objects (Müller et al., 2009, 2011), our results suggest that PEs to violations of specific features are processed mostly in different regions. Clusters in the cerebellum, thalamus, precuneus, posterior medial frontal cortex, and right temporal areas were activated exclusively for predictions and/or pwPEs pertaining to emotions. However, activations in the fusiform gyrus for color and emotion showed some overlap (Fig. 5A). In addition, we could not find any evidence for an interaction between PEs for different features when they are task-irrelevant and unattended. However, we only considered an interaction between color and emotion PEs at the level of the GLM and did not investigate possible interactions at the level of the perceptual model of the HGF. This would require establishing a novel HGF that incorporates interactions between hidden beliefs.

### Color PEs

Color processing involves the ventral visual pathway (Mesulam, 1998; Bartels and Zeki, 2000), where fMRI studies have shown strong color-related activations (Brewer et al., 2005; Solomon and Lennie, 2007; Barbur and Spang, 2008; Brouwer and Heeger, 2009). The location of the fusiform activation in our experiment is in agreement with “color-biased” regions in the ventral occipito-temporal cortex (Lafer-Sousa et al., 2016). To our knowledge, there have been no previous investigations of color processing from a PC-related perspective. Our results suggest the importance of pwPEs, as a putative signature of PC, for color perception.

### Emotion PEs and predictions

Facial emotions are non-verbal acts of communication that express emotional states and intentions, and are fundamental in social interactions (Fridlund, 1994; Frith, 2009). The social environment is not constant, and detecting changes in the emotional valence of facial expressions in our social space is important for socially successful behavior. Prior ERP studies (Susac et al., 2004; Kimura et al., 2012; Li et al., 2012; Csukly et al., 2013; Stefanics et al., 2012, 2018a; Astikainen et al., 2013; Fujimura and Okanoya, 2013; Xu et al., 2018) suggest that emotional expressions are processed in a few hundred milliseconds and stored in predictive memory representations. We found emotion pwPEs in a set of areas including the bilateral cerebellum, precuneus, thalamus, right lingual and supramarginal gyrus, as well as right posterior medial frontal cortex. We observed activations pertaining to emotion predictions in bilateral fusiform and the right superior temporal gyrus. Details of significant clusters are provided in Table 1. This pattern of results (Fig. 5A) is in line with the notion that emotion processing involves a mosaic-like set of affective, motor-related and sensory components (Bastiaansen et al., 2009). More specifically, it demonstrates pwPE/prediction activations in areas that previous work identified as activated by the processing of emotional faces (Fusar-Poli et al., 2009; E et al., 2014; Adamaszek et al., 2017) and theory of mind tasks, in particular the Mind in the Eyes task (Schurz et al., 2014).

In our current study we observed positive and negative betas for color and emotion pwPEs, respectively, which might reflect complementary neural mechanisms for predictive processing across distinct features. The notion that predictive coding across features can be mediated by qualitatively different mechanisms (Auksztulewicz et al., 2018) suggests domain-specific predictive signaling. As fMRI does not allow to measure detailed neural firing but rather represents the bulk signal of excitation and inhibition within a region (Logothetis, 2008), we cannot draw conclusions about specific mechanisms that could lead to this difference in the PE signal.

In summary, our findings demonstrate that the same physical stimulus can elicits separate feature-specific pwPE/prediction responses, depending on distinct predictions about its various attributes. This is in agreement with PC theories of perception. In future extensions of this work, models of effective connectivity could examine the signaling of pwPEs/predictions in cortical networks as postulated by PC.

## Acknowledgements

We acknowledge support by the University of Zurich (KES), the René and Susanne Braginsky Foundation (KES), and the Clinical Research Priority Program “Multiple Sclerosis” (GS, KES).

